# Computational model of chimeric antigen receptors explains site-specific phosphorylation kinetics

**DOI:** 10.1101/262527

**Authors:** Jennifer A. Rohrs, Dongqing Zheng, Nicholas A. Graham, Pin Wang, Stacey D. Finley

## Abstract

Chimeric antigen receptors (CARs) have recently been approved for the treatment of hematological malignancies, but our lack of understanding of the basic mechanisms that activate these proteins has made it difficult to optimize and control CAR-based therapies. In this study, we use phospho-proteomic mass spectrometry and mechanistic computational modeling to quantify the *in vitro* kinetics of individual tyrosine phosphorylation on a variety of CARs. We show that each of the ten tyrosine sites on the CD28-CD3ζ CAR is phosphorylated by LCK with distinct kinetics. The addition of CD28 at the N-terminal of CD3ζ increases the overall rate of CD3ζ phosphorylation. Our computational model identifies that LCK phosphorylates CD3ζ through a mechanism of competitive inhibition. This model agrees with previously published data in the literature and predicts that phosphatases in this system interact with CD3ζ through a similar mechanism of competitive inhibition. This quantitative modeling framework can be used to better understand CAR signaling and T cell activation.

## Introduction

One of most widely used methods for engineering a patient’s T cells to fight cancer is through the expression of chimeric antigen receptors (CARs). CARs are proteins that combine an extracellular antibody-derived targeting domain with intracellular T cell activating domains derived from the endogenous T cell receptor (1). These engineered T cells have emerged as promising treatments for hematopoietic cancers (2,3); however, not all patients respond to treatment and it has been difficult to expand these therapies to solid tumors (4–7). Significantly, it has been shown that CARs are less effective at activating T cells than engineered T cell receptors (TCRs) (8). More work needs to be done to better understand the mechanisms through which CAR-engineered T cells become activated so that they can be more optimally designed and expanded to a wider patient population. In this study, we use quantitative phospho-proteomic mass spectrometry and computational modeling to explore the mechanisms that lead to the phosphorylation of CAR proteins. Computational models, like the one developed here, provide a unique method to use basic engineering principles to better understand and optimize the signaling pathways that activate CAR-engineered T cells.

The CAR-T cell therapy Yescarta was approved by the FDA in October 2017 and contains signaling domains derived from the CD3ζ domain of the T cell receptor (TCR) and the CD28 co-stimulatory domain (3). These T cell signaling domains are phosphorylated by the Src family kinases, the most important of which in endogenous T cells is lymphocyte-specific protein tyrosine kinase (LCK) (9–11). CD3ζ contains six tyrosine phosphorylation sites, arranged in pairs on three immunoreceptor tyrosine-based activation motifs (ITAMs) (**Figure 1A**) (12). When doubly phosphorylated, these ITAMs can bind to the adaptor protein ZAP-70, perpetuating downstream signaling and also protecting the CD3ζ ITAMs from dephosphorylation (13,14). Importantly, in addition to this main form of activation through ZAP-70, the three ITAMs can also bind other signaling proteins. Literature data indicates that the ITAMs have different binding capabilities and, therefore, can induce different downstream signaling events (15); however, more work needs to be done to specifically enumerate how the individual ITAMs differ in both their activation and, subsequently, their downstream signaling.

Previous studies have attempted to qualitatively define the order of CD3ζ phosphorylation using CD3ζ phospho-tyrosine specific antibodies (16); but, the similarity between ITAM sites limited the antibody specificity, preventing reliable conclusions regarding the phosphorylation order. In 2003, Housden *et al.* used synthetic peptides that each contained one ITAM tyrosine to measure the preference of LCK for each site through radioactive phosphate incorporation. While they did find differences in the rates of tyrosine peptide phosphorylation, their experiments were performed in solution and each single tyrosine-bearing peptide was phosphorylated independently. Therefore, this study did not account for conformational, steric, and competitive factors that may influence the phosphorylation rates at different sites.

A study from Mukhopadhyay and colleagues proposed a mechanism for CD3ζ phosphorylation and dephosphorylation based on experimental measurements of total protein phosphorylation for CD3ζ with individual ITAM mutations (17,18). In this work, wild type or ITAM mutant CD3ζ was co-expressed with LCK and phosphatase CD148 in HEK293 cells. Total CD3ζ phosphorylation was measured as a function of phosphatase inhibition, and it was found that there is no significant difference between the individual ITAMs; however, increasing the number of ITAMs decreased the EC50 value of the phosphorylation curve without changing the Hill coefficient. Mukhopadhyay and colleagues hypothesized that the addition of phosphate groups on the CD3ζ intracellular domain could increase the rigidity of the CD3ζ tertiary structure, making unphosphorylated sites more accessible. They modeled this as an exponential increase in the association rate of the kinase and phosphatase toward CD3ζ; however, this mechanism is not fully validated without site-specific phosphorylation data.

Combining the CD3ζ activating domain and a co-stimulatory domain on the same protein adds additional complexity to the CAR. The CD28 co-stimulatory domain has four tyrosine sites, which can be phosphorylated by LCK (19), and may also influence the catalytic activity of LCK (20,21). Once phosphorylated, CD28 tyrosine sites bind to various adaptor proteins that are also phosphorylated downstream of CD3ζ (22). Thus, CD28 can tune the response to CD3ζ activation. Additionally, the recruitment and competition by CD28 for LCK may alter the phosphorylation of CD3ζ.

All ten of the tyrosine sites on CD3ζ and CD28 work together, in different ways, to affect the downstream signaling that controls T cell activation responses such as cytotoxicity, cytokine production, proliferation, and survival (23–27). By better understanding how these chimeric proteins are phosphorylated, we can identify ways to tune them to create more optimal CAR therapies. In this study, we have explored the kinetics of CD3ζ and CD28 phosphorylation in detail. We paired a recombinant protein system with phospho-proteomic mass spectrometry to measure the site-specific phosphorylation of CAR proteins by LCK over time. To our knowledge, this is the first study to quantify phosphorylation at individual sites on the intact intracellular domain of a CAR protein. We then fit this data using a computational model to robustly quantify the differences between the phosphorylation kinetics of the ten tyrosine sites. We used the computational model to generate new predictions regarding the mechanism with which LCK phosphorylates the CD3ζ ITAMs. We can use the novel insights from this study to continue expanding our understanding of CAR-mediated T cell activation and better engineer future CAR therapies.

## Methods

### Recombinant protein expression and purification

His_10_-KKCK-CD3ζ in the pET28a vector and HIS_10_-LCK-G2A in the pFastBacHTA vector were a kind gift from Dr. Ronald Vale (28). To make the HIS_10_-CD28-CD3ζ recombinant protein, the DNA sequence for the intracellular domain of CD28 (aa 180-220) was codon optimized and constructed by Integrated DNA Technologies (IDT-DNA). This sequence was then cloned directly upstream of CD3ζ in the pET28a vector using Gibson assembly. All individual point mutations in the CAR vectors were introduced by the QuikChange XL site-directed mutagenesis kit (Agilent).

The sequence for HIS_10_-LCK-G2A was amplified out of the pFastBacHTA vector by PCR and cloned into the FUW vector through Gibson assembly. LCK is able to undergo autophosphorylation at both activating and inhibitory tyrosine sites (Y394 and Y505, respectively). We previously showed that when LCK is phosphorylated at these tyrosine residues, it has differential catalytic activity (29). Therefore, to exclude any confounding effects due to changes in enzymatic efficiency, we used a constitutively active form of LCK containing a tyrosine to phenylalanine mutation at the inhibitory site (Y505F). This point mutation was introduced by the QuikChange XL site-directed mutagenesis kit (Agilent).

All CAR proteins were expressed in the BL28(DE3) strain of E. coli cells. E. coli cells were lysed as described in (30). His_10_-LCK-G2A-Y505F was transiently expressed in HEK293T cells through a standard calcium phosphate precipitation protocol (31). 48 hours after transfection, HEK293T cells were lysed in buffer containing 20 mM TrisHCl, pH 7.5, 600 mM NaCl, 2 mM MgCl2, 5 mM immidazole, 10% glycerol, 1% NP-40, 1 mM Na3VO4, 10 mM NaF, and 1× complete protease inhibitor (Roche). All HIS_10_ proteins were purified using FPLC, first on a Ni-NTA agarose column followed by gel purification using the HiPrep 16/60 Sephacryl S-200 HR column (GE Life Sciences) in HEPES-buffered saline (HBS) solution containing 50 mM HEPES-NaOH (pH 7.5), 150 mM NaCl, and 10% glycerol, as described in (28). Protein monomer fractions were concentrated, snap frozen in liquid nitrogen, and stored at −80°C. All purified recombinant proteins were quantified by SDS-PAGE and Coomassie staining using BSA as a standard.

Mass spectrometry confirmed that nearly 100% of the purified LCK-Y505 is phosphorylated at the activating Y394 site, while 100% of the CAR proteins were unphosphorylated after purification.

### Liposome preparation

Synthetic 1,2-dioleoyl-sn-glycero-3-phosphocholine (POPC), 1-palmitoyl-2-oleoyl-sn-glycero-3-phospho-l-serine (POPS), and 1,2-dioleoyl-sn-glycero-3-[(N-(5-amino-1- carboxypentyl)iminodiacetic acid)succinyl] (nickel salt, DGS-NTA-Ni) were purchased from Avanti Polar Lipids and resuspended in chloroform. Liposomes were prepared as described in (28). Briefly, phospholipids (80% POPC, 10% POPS, 10% DGS-NTA-Ni) were dried as thin films under Ar gas and desiccated overnight. The lipids were then resuspended in 1x kinase buffer (50 mM HEPES-NaOH (pH 7.5), 150 mM NaCl, 10 mM MgCl2, 1 mM TCEP), and subjected to 5x freeze thaw cycles. The lipid mixture was then extruded through 200 nm pore-size polycarbonate filters to produce large unilamellar liposomes. As such, we assume that the liposomes have an outer diameter of roughly 200 nm (28). For liposomes with varying POPS concentration, the amount was compensated for by adjusting the POPC concentration.

### Protein phosphorylation time courses

His-tagged LCK and CAR proteins were mixed with Ni-bearing liposomes for 1 hour to allow for the proteins to attach to nickel bearing lipids on the surface of the liposome, as calculated and described in (28). We used 20,000 molecules/μm^2^ CAR proteins and titrated down LCK to a very low concentration to allow us to distinguish the differences between the CD3ζ site phosphorylation kinetics. The low LCK concentration allows us to measure the rapid phosphorylation kinetics of CAR tyrosine sites. Estimation of the final LCK concentration is described in the *Mechanistic computational modeling* section. Once the proteins attached to the liposomes, 10x ATP in kinase buffer was added to a final concentration of 1 M. Samples were taken at various times and the reaction was | stopped by adding urea to 8_M and boiling for 5 minutes. Time samples were then frozen at −20°C until they were prepared for phospho-proteomic mass spectrometry.

### Standard curve preparation

Because differing ionization efficiencies for each peptide result in different intensities for the same amount of peptide on a mass spectrometer (MS), a standard curve is necessary to compare between peptide MS intensities in a sample. We constructed our standard curves based on a known ratio of phosphorylated:unphosphorylated peptide. For each CAR protein, we quantified the amount of protein and aliquoted the same volume from a given sample into two vials. To one vial we added LCK and ATP and let LCK phosphorylate the CAR overnight at room temperature. To the other vial, we added equal volumes of HBS buffer, so that the phosphorylated and unphosphorylated CAR proteins would remain at the same concentration. The next morning, urea was added to both vials to a final concentration of 8 M. We then combined various volume ratios of the two solutions to create a standard curve with known ratios of phosphorylated:unphosphorylated peptides. The standard curve samples were stored at −20°C until they were ready to be prepared for analysis by mass spec.

### Phospho-proteomic sample preparation

The time course and standard curve samples were thawed to room temperature and reduced by the addition of DTT to a final concentration of 5 mM for 1 hr at 37°C. Samples were next alkylated with iodoacetamide at a final concentration of 25 mM for 1 hr at room temperature in the dark. This reaction was quenched by the addition of DTT to a final concentration of 10 mM for 30 min. Samples were then diluted to a final urea concentration of 2 M with 100 mM Tris, pH 8, and trypsin digested overnight at 37°C. The next morning, samples were acidified to a pH<4 with 5% TFA, purified by C18 zip-tip (Millipore) according to the manufacturer’s instructions, and eluted into 50% acetonitrile solution. Purified samples were then dried and stored at −80°C until they were ready to be analyzed.

### Phospho-proteomic data collection

All data samples were run in technical duplicates. Data was collected in three sets, each with their own standard curve, shown in **Supplemental Figure S1**: (1) first biological replicate of the wild type ITAM phosphorylation on 10% POPS liposomes, the wild type ITAM phosphorylation on 10% and 45% POPS liposomes, and a CD3ζ standard curve, (2) the second biological replicate of the wild type ITAM phosphorylation on 10% POPS liposomes, the individual tyrosine to phenylalanine CD3ζ ITAM point mutants, and a CD3ζ standard curve, (3) all 28ζ proteins, including the Y206F and Y209F mutants, and a standard curve for 28ζ, CD28-Y206F-CD3ζ, and CD28-Y209F-CD3ζ. Desalted samples were reconstituted in buffer A (0.1% formic acid). The samples were injected into an Easy 1200 nanoLC ultra-high pressure liquid chromatography coupled to a Q-Exactive Plus mass spectrometer (Thermo Fisher Scientific). Peptides were separated by reversed-phase chromatography (PepMap RSLC C18, 2μm, 100Å, 75 μm X 15 cm). The flow rate was set to 300 nl/min at a gradient starting with 6% buffer B (0.1% FA, 80% acetonitrile) to 55% B in 25 minutes, followed by an 8 minutes washing step to 100% B. The maximum pressure was set to 1180 bar and column temperature was held constant at 50 °C.

Peptides separated by the column were ionized at 2.0 kV in positive mode. MS1 survey scans were acquired at resolution of 70k from 275 to 1500 m/z, with maximum injection time of 80 ms and AGC target of 1e6. MS/MS fragmentation of the 10 most abundant ions were analyzed at a resolution of 35k, AGC target 1e5, maximum injection time 100 ms, and normalized collision energy 25. Dynamic exclusion was set to 10 s and ions with charge 1, 7, and >7 were excluded.

### Mass spec data analysis and normalization

MS/MS fragmentation spectra were searched with Proteome Discoverer SEQUEST (version 1.4, Thermo Scientific) against the recombinant protein sequences (17 entries) used in this study. The maximum missed cleavages was set to 2. Dynamic modifications were set to oxidation on methionine, phosphorylation on serine, threonine, and tyrosine, and acetylation on protein N-terminus. Fixed modification was set to carbamidomethylation on cysteine residues. The maximum parental mass error was set to 10 ppm and the MS/MS mass tolerance was set to 0.02 Da. False Discovery threshold was set to 0.01 using Percolator node validated by q-value. Phosphosite localization was confirmed using PhosphoRS (all sites >75% probability).

MS1 peak quantification was performed manually in Skyline (version 3.7) for each phosphorylated/unphosphorylated peptide pair. We analyzed only peptides with no missed cleavages and no modifications other than tyrosine phosphorylation, which were consistently the largest peaks. One exception was made for the CD28 peptide containing tyrosine site Y218. The unphosphorylated form of this peptide was smaller than the cutoff mass to charge ratio used in our data collection. Therefore, we analyzed this site using the peptide with one N-terminal missed cleavage.

To create our peptide standard curves, we calculated the ratio of each phosphorylated/unphosphorylated peptide, plotted them against the known ratios and fit the resulting linear plots (**Supplemental Figure S1**). Technical replicates of each peptide were combined together to fit the standard curves so that one standard curve was used to normalize the phosphorylated/unphosphorylated peptide intensity ratios for each set of peptide time course technical replicates. We then used the normalized ratios to calculate the percent phosphorylation over time for each time course technical replicate and used the two sets to calculate the mean and standard deviation of the data. Time courses were only normalized to the standard curve data collected at the same time.

### Statistical analysis

All statistical analyses were done using a one-way ANOVA followed by multiple pairwise comparisons using the Tukey t-test in Prism (version 7, GraphPad).

### Sigmoidal parameter calculations

Data was fit in Prism (version 7, GraphPad) to a standard sigmoidal curve with plateaus at 0 and 100%.

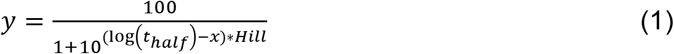

Where *x* is the time on a log scale, *y* is the output, *t*_*half*_ is the half maximal time, and *Hill* is the Hill coefficient. For the comparison of *t*_*half*_ and *Hill* for the random and sequential models, the models were first fit to data using MATLAB, as described in the “Mechanistic computational modeling” section below. The model responses were then entered into Prism as data sets and fit to Equation 1.

### Model implementations

For the sequential and random order models, the ordinary differential equations were written using standard Michaelis-Menten kinetics, as shown in Equation 2.

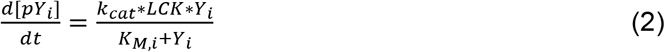

Where *Y*_*i*_ and *pY*_*i*_ represent the unphosphorylated and phosphorylated species, respectively, for ITAM tyrosine site *i*. Here, *i* can be A1, A2, B1, B2, C1 or C2. *LCK* represents the concentration of the kinase LCK, *k*_*cat*_ is the catalytic rate, and *K_M,i_* is the Michaelis-Menten constant for each ITAM site *i*.

For the phosphate dependent model, we also used random order Michaelis-Menten kinetics. However, the Michaelis-Menten constant for each ITAM site, *K_M,i_*, was scaled by a constant, *λ*, raised to the power of the number of phosphate groups on the indicated CD3ζ molecule, *p*, resulting in, 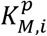, as shown in Equation (3).

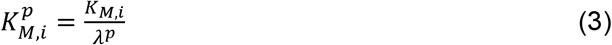

We started with a *λ* value of three, as estimated by Mukhopadhyay et al. (17), and fit the *k*_*cat*_, six *K_M,i_*values, and the total amount of LCK kinase to the data. We then expanded our parameter space to explore other values of *λ*. We found that *λ* values less than one were able to significantly improve the fit. However, this inversion of the *λ* parameter resulted in a mechanism that was deemed physiologically irrelevant based on previous work in the literature (17); therefore, *λ* was kept at a value of three.

The competitive inhibition model also relied upon Michaelis-Menten kinetics. The equation describing the rate of each ITAM phosphorylation is shown in Equation 4.

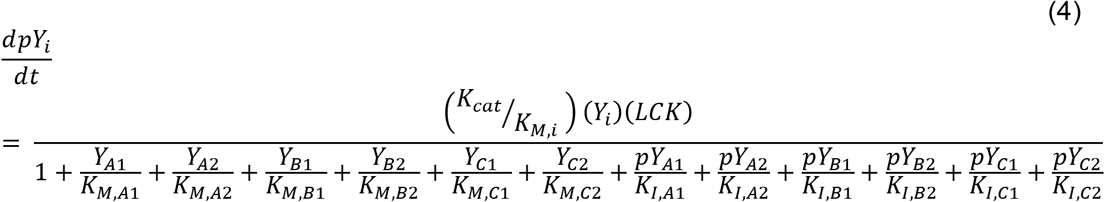

Where all variables are the same as described for Equation 1, and *K_l,i_* is the inhibitory constant for each ITAM site, *i*.

In our preliminary exploration of the model parameter space, we identified several groups of parameters in this model structure that were correlated: the Michaelis-Menten constants, the inhibition constants, and the total LCK concentration and catalytic rate. Therefore, to better constrain this system, we made a series of assumptions. First, the addition of six inhibition constants greatly over parameterizes the model. To reduce this number, we assume that adding a phosphate group will affect all of the ITAM sites the same way. Therefore, we simplified the inhibition constants to a single factor (*X*_*l*_) that could be used to scale the Michaelis-Menten constant for each site, as shown in Equation 5.

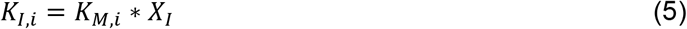

Second, we fixed the catalytic rate of LCK. The initial conditions for the CAR proteins were based on the measured protein densities used in our experiments, 20,000 molecules/μm^2^. In order to be able to distinguish between the rapid phosphorylation kinetics of CAR tyrosine sites, low concentrations of LCK were used in the experiments. This experimental condition also agrees with the assumption of Michaelis-Menten kinetics that the enzyme concentration is much less than the substrate concentration. However, this low level of LCK made it very difficult to measure the exact concentration relative to the experimental errors. Therefore, we held the catalytic rate constant based on the average rate of CD3ζ phosphorylation by constitutively active LCK, calculated in (28) and fit the initial concentration of LCK.

Third, as all of the *K*_*M*_ values varied together in a correlated manner, we chose to hold the *K*_*M*_ value for site B1, *K_M,B1_*, which is phosphorylated at an intermediate rate relative to the other sites, equal to the estimated *K*_*M*_ value from (28). Through many simulations, we confirmed that these parameter assumptions did not significantly affect the model fit to the data.

### Comparison of model structures

Our mechanistic computational models were written as a set of rules in BioNetGen (32) and implemented in MATLAB (**Supplemental Files S1-S5**). We used Michaelis-Menten kinetics to describe the reaction rate of LCK toward each of the six ITAM sites, as described in the previous section. Parameter fitting was performed in an iterative manner. For the different model structures, starting values for Michaelis-Menten constants were first identified by manually performing parameter sweeps across a wide range. For this step, the catalytic rate was held at 360 min^−1^, based on literature values (28), and total LCK concentration was 60 molecules/μm^2^, as estimated in the experimental setup. Once a suitable order of magnitude was identified for the K_M_ values, all of the parameters were allowed to vary two-fold up and down, and the parameters were fit to all of the data together (wild type CD3ζ, mutant CD3ζ, and liposome concentration data) using particle swarm optimization (PSO) (33). Each data set was fit a minimum of 10 times. The error between the model fits and experimental data (calculated as the sum of the squared residual) was used to characterize the goodness of fit.

For model structures that did not fit the data well, we further explored the parameter space to see if changing the range of parameters could improve the fit. To do this, we manually altered the parameters for which the physiological range is not well defined (i.e. the two-dimensional Michaelis-Menten constants, and any scaling factors). If we identified a parameter space that better represented the data, we performed another round of 10 parameter set fits, using PSO. For the final parameter estimation, all of the parameters that were not fixed were allowed to vary two orders of magnitude up and down from their baseline values (*LCK* = 60 molecules/μm^2^, *K_M,i_* = 270 molecules/μm^2^, and *X*_*l*_ = 1). The model was fit 100 times using PSO and the 50 best fits were taken as the final parameter ranges. For all models, the parameter set with the best fit was used for model comparison.

### Phosphatase model

To explore the mechanism of phosphatase activity on the model, we implemented phosphatase mechanisms with a random order, a phosphate dependent mechanism, or a competitive inhibition mechanism. For the phosphate dependent model, we held the phosphate dependent scaling factor, *λ*, equal to three, as estimated by Mukhopadhyay et al. In the competitive inhibition model, both phosphorylated and unphosphorylated ITAM sites provide competitive inhibition. For each of these mechanisms we manually explored ranges of parameter space within one order of magnitude above or below the value that was estimated for the LCK phosphorylation parameters. We particularly focused on variations in the parameters for which all of the phosphatase Michaelis-Menten kinetics were (i) all the same between the ITAM sites, (ii) scaled so that they maintained the same relative differences as was identified for the LCK parameters. Once a mechanism and parameter space with the correct trends for EC50 and Hill coefficients were identified, we then further tuned the phosphatase Michaelis-Menten constants to better fit the data.

## Results

### The six tyrosine sites on CD3ζ are phosphorylated by LCK with different kinetics

We first sought to explore how LCK phosphorylates the six tyrosine sites on CD3ζ. To do this, we utilized a liposome-based recombinant protein system, developed by Hui and Vale (28). In this system, His-tagged proteins are bound to nickel chelating lipids on the surface of large unilamellar liposomes, as shown in **Figure 1A**. Since the CAR and LCK proteins are largely membrane bound in T cells, this system allows us to mimic the two-dimensional protein arrangement and more accurately capture the true kinetics of the interactions between these proteins.

**Figure 1:**
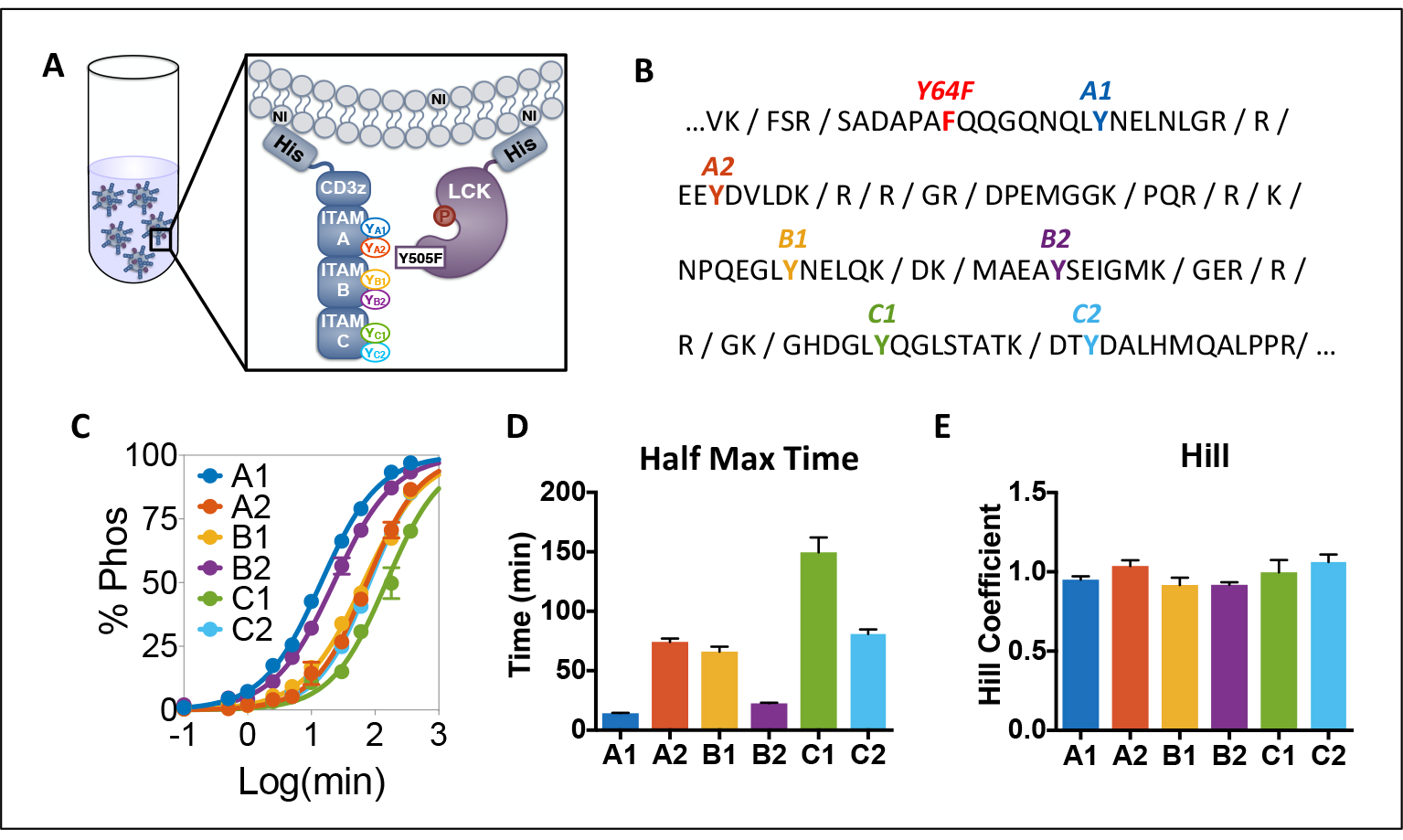
CD3ζ sites are phosphorylated by LCK with different kinetics. (A) Schematic of the experimental liposomal system. CD3ζ and LCK His-tagged proteins were purified and allowed to bind to large unilamellar liposomes bearing nickel-chelated lipids. Once proteins were bound, ATP was added, and the proteins were allowed to interact for various times before being subjected to phospho-proteomic mass spectrometry for quantification. (B) Sequence of CD3ζ intracellular domain with trypsin cut sites denoted. Individual ITAM tyrosine sites are labeled in different colors. Y64F indicates a tyrosine to phenylalanine mutation to ensure that each peptide only has one tyrosine phosphorylation site. This mutation does not influence overall phosphorylation kinetics (see Supplemental Figure S2). (C) Experimental data (circles) and sigmoidal fit (lines) for CD3ζ ITAM phosphorylation on liposomes containing 10% acidic POPS lipids. Error bars represent the standard deviation of two technical replicates normalized by site-specific standard curves. (D) Half maximal time for each CD3ζ ITAM site. Data represents mean and standard error of the mean (SEM) of the fit to a 4-parameter sigmoidal curve. (E) Hill coefficient for each CD3ζ ITAM site. Data represents mean and SEM of the fit to a 4-parameter sigmoidal curve.

The liposome-bound proteins were allowed to react in the presence of ATP, and we performed phospho-proteomic mass spectrometry to specifically measure the phosphorylation at each ITAM site over time. To quantify the site-specific phosphorylation, we needed to directly compare the mass spectrometry intensity of phosphorylated and unphosphorylated peptide pairs. To do this, we used a standard curve with a known ratio of phosphorylated:unphosphorylated peptide (**Supplemental Figure S1**) (34). Additionally, we needed to ensure that there is only one tyrosine site on each tryptic peptide. This is true for all CD3ζ ITAM tyrosine sites except A1 (**Figure 1B**). The peptide containing site A1 also contains a tyrosine at position 64, which is not part of an ITAM and is not predicted to play a significant role in CD3ζ phosphorylation based on known LCK binding motifs and computational predictions (35–38). Therefore, we added a Y64F mutation in the CD3ζ recombinant protein and validated that it does not influence the overall phosphorylation kinetics within this system (**Supplemental Figure S2**). In this way, we were able to normalize the phosphorylated:unphosphorylated intensity ratios for each ITAM site in our time courses by the standard curves, thus calculating the percent phosphorylation over time for each tyrosine site of interest.

**Figure 1C** (**dots**) shows the percent phosphorylation of each of the six ITAM sites over time on liposomes that contain 10% acidic phosphatidylserine (POPS) lipids, which is similar to the concentration of phosphatidylserine on the inner leaflet of the T cell plasma membrane (28). Our measurements show that the sites are not phosphorylated at the same rate. To quantify the differences, we fit these data to a four-parameter sigmoidal curve (**Figure 1C, lines**), estimating the half maximal time (**Figure 1D**) and the Hill coefficient (**Figure 1E**) for each site. The half maximal times show that the six sites are phosphorylated with different kinetics (A1>B2>B1≥A2≥C2>C1). In comparison, the Hill coefficients for all tyrosine phosphorylation sites are close to one.

### CD3ζ ITAM mutations

We next wanted to explore the influence that individual tyrosine sites have over the phosphorylated kinetics of other sites. Specifically, we wanted to identify if there are any binding or competitive effects that influence the kinetics at distant sites. Therefore, we individually mutated each tyrosine site to a phenylalanine and measured the percent phosphorylation of the other sites over time.

We also investigated the effect of the liposome membrane acidity. Several T cell receptor proteins, including the closely related CD3ε and CD28 proteins, have been shown to have basic residues in their intracellular domains that can interact with acidic lipids on the inner leaflet of the T cell membrane (39–41). These interactions are thought to help limit tyrosine accessibility, thus controlling aberrant phosphorylation in unstimulated cells. Therefore, we also tested if CD3ζ interactions with the acidic POPS lipids in the liposome membrane were contributing to the different rates of phosphorylation seen in the site-specific data.

**Figure 2** shows the overlay of all of the phosphorylation time course experiments (six individual CD3ζ Y to F point mutations, wild type CD3ζ stimulated on 0% and 45% POPS liposomes, and two biological replicates of wild type CD3ζ stimulated on 10% POPS liposomes) for each site. Although there is some variability in the phosphorylation time courses between the mutations, the trends for all of the site-specific time courses are very similar. Additionally, individual site mutations and changes to the acidic lipid microenvironment do not significantly affect the order of the phosphorylation kinetics of CD3ζ tyrosine sites.

**Figure 2:**
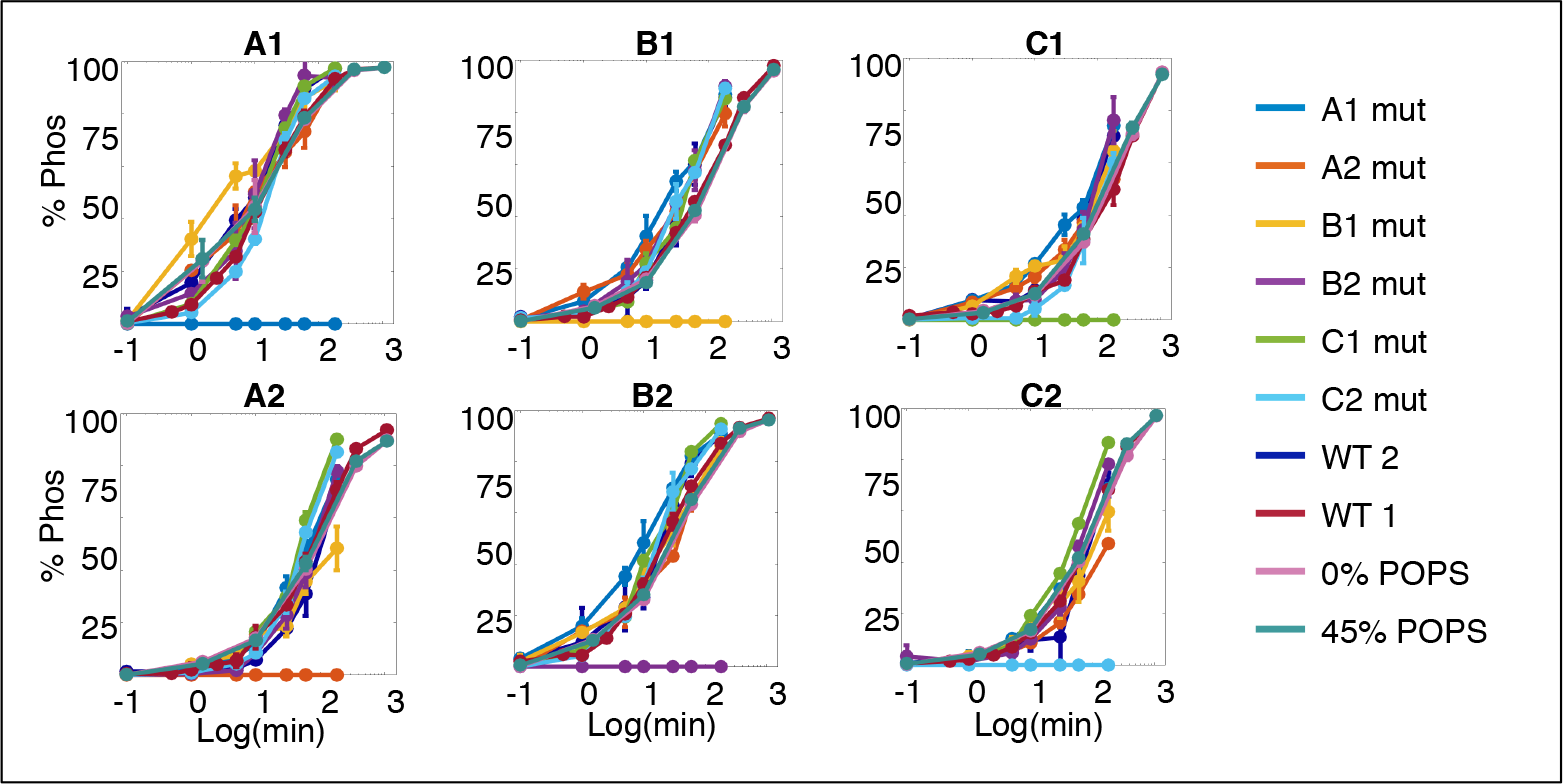
Comparison of individual tyrosine site mutations on phosphorylation kinetics. Experimental data for each CD3ζ ITAM site for different experimental conditions: WT 1 and 2 - biological replicates of CD3ζ with unmutated ITAMs on liposomes containing 10% POPS, XX mut (where XX represents the tyrosine to phenylalanine ITAM mutation site for CD3ζ stimulated on 10% POPS liposomes), and X% PS (where X represents the POPS concentration for liposomes bearing CD3ζ with wild type ITAMs). Error bars represent the standard deviation of two technical replicates normalized by site-specific standard curves.

To further compare between the sites, we grouped the time course responses together for all of the experimental conditions and used the pair-wise Tukey t-test to identify which ITAM site phosphorylation levels were significantly different from the others (**Supplemental Figure S3**). We compared the data sets at two different time points 10 minutes (blue), which is close to the half maximal time of the quickly phosphorylated ITAM sites A1 and B2, and 60 minutes (orange), which is close to the half maximal time of the majority of the other sites. The 10-minute comparison shows that A2, C1, and C2 are not significantly different from each other, while site B1 is significantly different from A2 and C1 but not C2. From both the 10-minute and 60-minute comparisons, we see that sites A1 and B2 are both significantly different from all other sites.

### Mechanism of CD3ζ phosphorylation by LCK

This data shows us key features of the kinetics of CD3ζ site-specific activation. To determine a specific mechanism of interaction between LCK and CD3ζ, we turned to computational mechanistic modeling. We explored a variety of different mechanisms described in the literature or indicated by the data itself (see Methods for further explanation). We fit each of these various models to all of our data (wild type CD3ζ, tyrosine to phenylalanine mutant CD3ζ, and changes to the liposome composition) and analyzed the results to make a hypothesis about which mechanism best represents the system. The model mechanisms were compared based on the overall fit to the data, as well as the half maximal time and Hill coefficient of each model predicted phosphorylation time course.

#### Sequential order

The first mechanism we tested was a sequential phosphorylation order, in which LCK can phosphorylate the six CD3ζ in a specified order defined by the order of the half maximal time from the sigmoidal fit to the data (**Figure 1D**). These reactions are modeled using Michaelis-Menten interactions (**best fit parameters are listed in the BioNetGen Supplemental File S1**). As shown in **Figure 3A**, this model is able to capture the differences in half maximal time well, but it leads to a consistent increase in the Hill coefficient for each subsequent tyrosine site in the sequence (A1, B2, B1, A2, C2, and C1). This increase in slope, referred to as ultrasensitivity, has been described as a characteristic of sequential multi-site phosphorylation previously (18,42,43). However, this ultrasensitivity does not match our raw data, which shows a consistent Hill coefficient for all sites (**Figure 1E**) or other data of CD3ζ phosphorylation in the literature (17). Additionally, the shape of the model fits does not qualitatively match the experimental data. Overall, the modeling results indicate that LCK does not phosphorylate CD3ζ sequentially.

**Figure 3:**
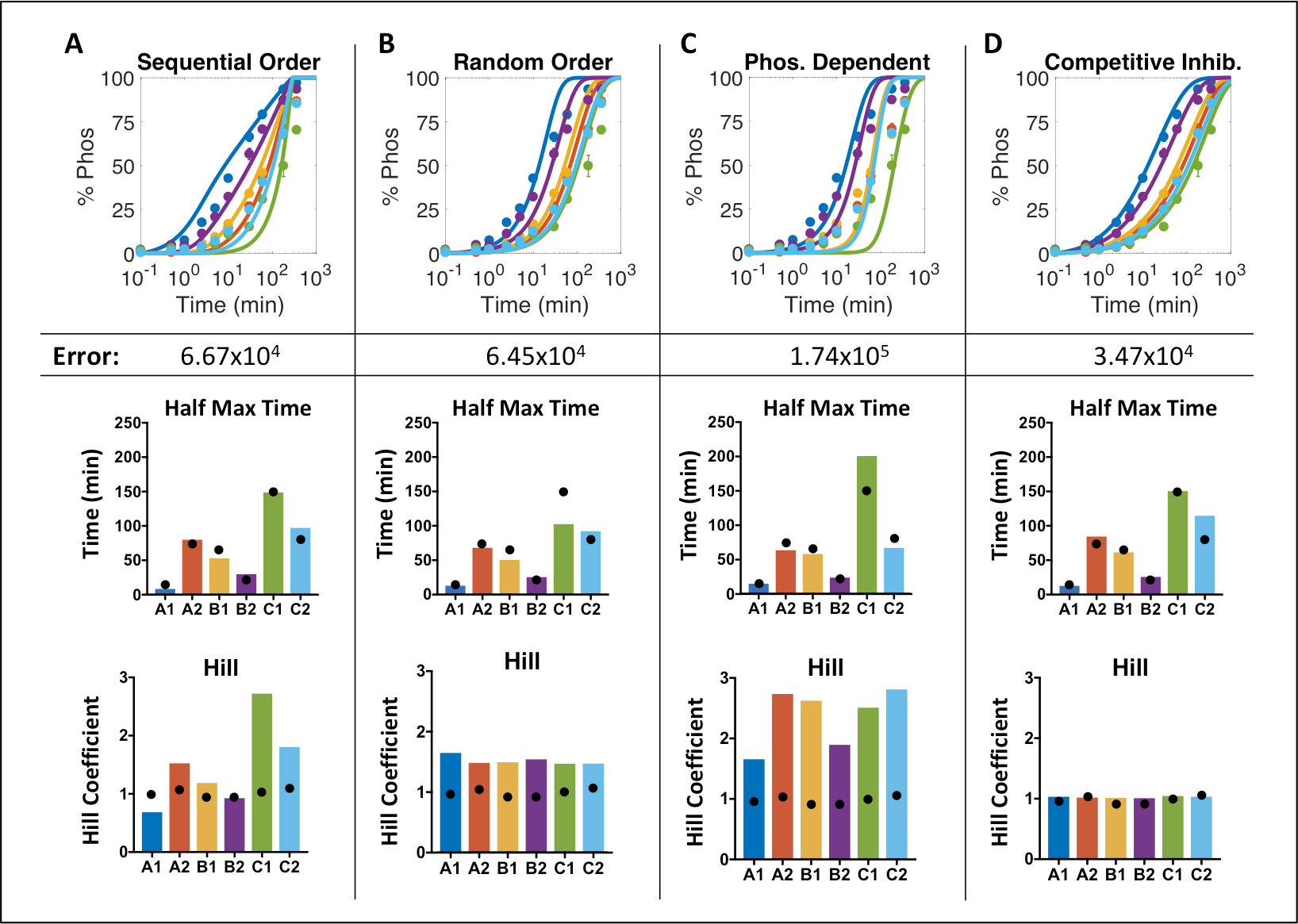
Comparison of CD3ζ phosphorylation mechanisms. Model analysis for mechanisms of (A) Sequential order phosphorylation, (B) Random order phosphorylation, (C) Phosphate dependent phosphorylation rates, and (D) Competitive inhibition by unphosphorylated and phosphorylated CD3ζ ITAM sites. (Top) Model fit to experimental data. Error represents the residual error between the model and the data for all data sets, including wild type, individual tyrosine to phenylalanine mutants, and different liposome concentrations. (Middle) Half maximal times of the model predictions. (Bottom) Hill coefficients of the model predictions. Black dots indicate the mean values from the sigmoidal fit to the data shown in Figure 1 D and F.

#### Random order

We next tried a simple mechanism of a random phosphorylation order. In this random order model, each of the six ITAM sites interact with LCK independently using Michaelis-Menten kinetics (**Figure 3B, best fit parameters are listed in the BioNetGen Supplemental File S2**). With this model structure, each of the sites have the same Hill coefficient, which is similar to the experimental data, although the model Hill coefficient is slightly higher than that estimated for the data itself. Additionally, the random order model has a lower residual error, and thus fits the data better than the sequential model. Upon visual inspection, we can see that the random order model better captures the overall trends of the data. However, it consistently underestimates the level of phosphorylation at early time points and overestimates the gradual approach to saturation seen at later time points in the data. Therefore, we continued to explore other, more complex, models of random phosphorylation to find a mechanism that could better represent our data.

#### Phosphate dependent

We, therefore, turned to a previously proposed mechanisms in the literature, one given by Mukhopadhyay and colleagues and described in the introduction. In this mechanism, the addition of phosphate groups to CD3ζ causes an increase in the accessibility of unphosphorylated sites (17). In this model, a phosphate dependent scaling factor, *λ*, is used to adjust the Michaelis-Menten constant for each ITAM site, *K_M,i_* (see methods). For phosphate groups to increase binding in our model, *λ*, must be greater than one. Using a constant *λ* equal to three, predicted by Mukhopadhyay and colleagues, the model is not able to fit our data well (**Figure 3C, best fit parameters are listed in the BioNetGen Supplemental File S3**). Although the predicted response is able to capture differences in the half maximal time between the sites, it does not fit the early or late phosphorylation time courses well. Additionally, the Hill coefficients are much higher than those of the data itself, and they show site-specific differences. This is because the sites that are phosphorylated later start with higher effective Michaelis-Menten constants due to the scaling from sites that are already phosphorylated. Upon a wider parameter search, we find that having a *λ* value less than one will result in a significantly better fit to the data (residual error = 3.72×10^4^), as this will increase the *K*_*M*_ value for high levels of CD3ζ phosphorylation, allowing for the system to slow down at later time points. However, this inversion of the *λ* value would contradict the biological hypothesis proposed by Mukhopadhyay and colleagues and is, therefore, not feasible.

#### Competitive inhibition

Since the previously described models in the literature do not accurately reflect our site-specific data, we sought to identify a new mechanism, which could more accurately fit our experimental data while still agreeing with published data, including work by Mukhopadhyay et al. Using the random order Michaelis-Menten model as a starting point, we modified the equations to address the overestimation at later time points. We implemented a mechanism of competitive inhibition, in which the unphosphorylated and phosphorylated tyrosine sites could interact to compete with each other. Competitive inhibition and product inhibition have been show to play a role in other systems with multiple phosphorylation sites (42,44), indicating that it may play a role in CD3ζ phosphorylation.

The competitive inhibition model is able to fit the data with the lowest error of all the mechanisms we explored (**Figure 3D, best fit parameters are listed in the BioNetGen Supplemental File S4**). Additionally, it is able to capture the differences between the half maximal phosphorylation times while maintaining the same Hill coefficient throughout the system, matching our experimental data (**Figure 1E**). We also implemented two additional variations of the competitive inhibition model, in which only phosphorylated or unphosphorylated CD3ζ tyrosine sites compete for LCK activity. The model that only allowed competitive inhibition between the unphosphorylated sites was not able to fit the data well (residual error = 1.32×10^5^). On the other hand, only having inhibition from the phosphorylated species was better able to fit the data (residual error = 4.02×10^4^), indicating that product inhibition plays a more significant role in this system. Ultimately, the best fit to the data was given by a mechanism including both competitive inhibition from the unphosphorylated ITAM sites and product inhibition from phosphorylated sites (residual error = 3.47×10^4^). While the effect of competitive inhibition from the unphosphorylated sites is less significant than that of the product, physiologically, since the sites are in such close proximity, it is clear that all of these sites are interacting with LCK together, and this model accounts for that interaction.

To further validate that the competitive inhibition model mechanism gives a better fit than the phosphate dependent model, we tested a model structure that combines both of these features. This model mechanism did not give a significantly better fit to the data than the competitive inhibition model alone. Additionally, the effect of the phosphatase inhibition was negligible compared to the competitive inhibition effect, as evidenced by the fact that increasing the value of *λ* two orders of magnitude did not significantly influence the model. Ultimately, we conclude that a model of competitive inhibition by both phosphorylated and unphosphorylated sites on CD3ζ is the mechanism that best represents the data.

To estimate a final set of physiologically relevant parameter values, the LCK catalytic rate and *K*_*M*_ value for ITAM site B1, *K_M,B1_*, were held constant based on values in the literature (28). Additionally, a single scaling factor was used to estimate the inhibitory constants such that all of the parameters estimated were identifiable (see methods). The final parameter distributions for the 50 best sets are shown in **Figure 4**. From the standard deviation, we can see that the model consistently estimates parameter values within a narrow range. The estimated *K*_*M*_values correspond directly with the differences in half maximal time between the individual sites. These various parameter sets are able to fit all of the data, including CD3ζ mutant and liposome concentration data sets, similarly well (**Supplemental Figure S4**), indicating that the slight variation in the phosphorylation rates due to the individual site mutations can be account for by the competitive inhibition between sites. The same fits can be achieved by estimating different catalytic rates for each ITAM site and keeping the same *K*_*M*_ values between the ITAM sites as long as the competitive inhibition is still present (results not shown). Altogether, the modeling results provide confidence that the mechanism of competitive inhibition described here can accurately reflect the way in which LCK is able to phosphorylate CD3ζ.

**Figure 4:**
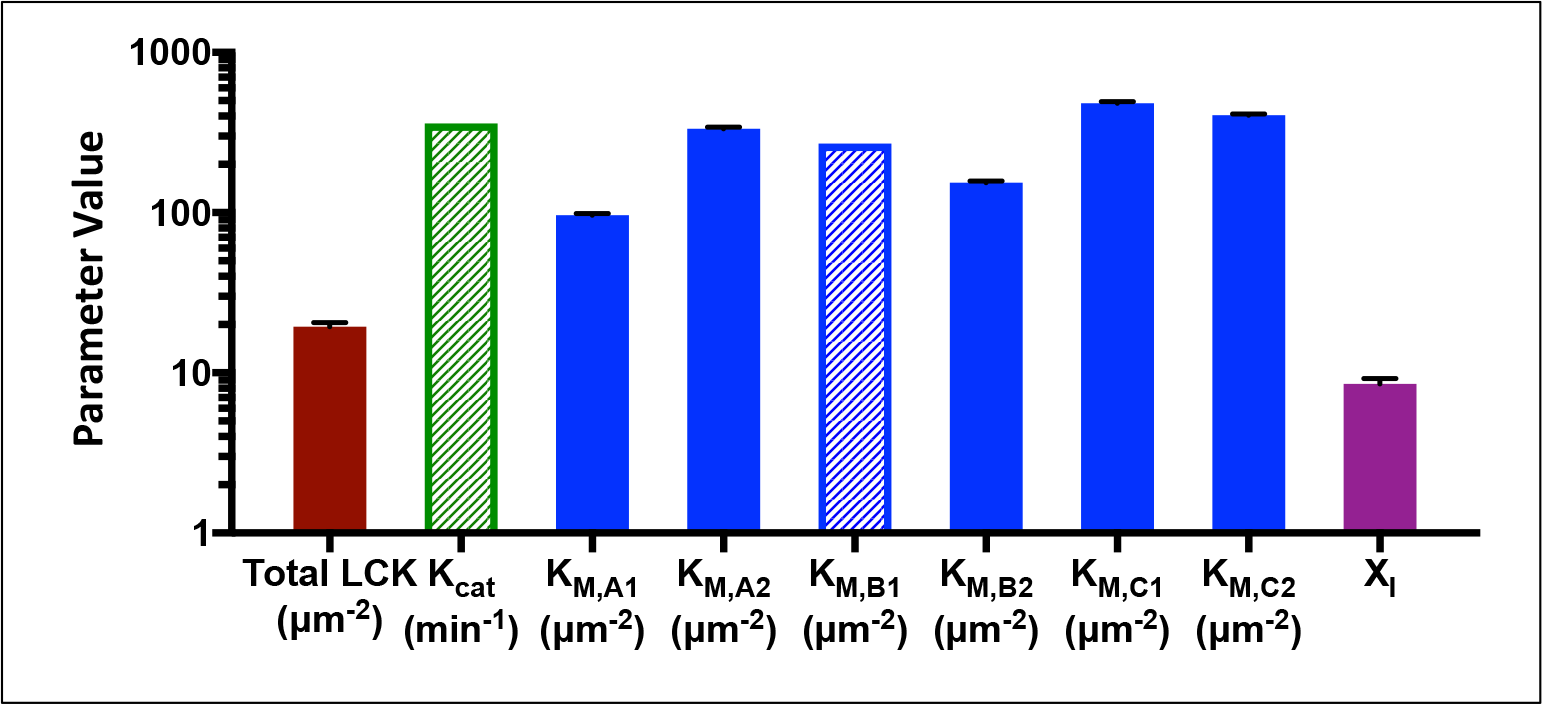
Estimated parameter sets. Solid bars show the mean and standard deviation of the 50 best fit parameter sets. Shaded bars show the value of parameters that were held constant during fitting.

### Effects of ITAM mutations can be explained by differences in LCK and phosphatase K_M_ parameter values

For this competitive inhibition mechanism of CD3ζ phosphorylation to be validated, it must be consistent with other experimental data of CD3ζ phosphorylation. To test this, we explored how this new model mechanism is able to reproduce experimental data from (17). Since the experimental data from that study indicated that there was no significant difference between the phosphorylation states of individual ITAMs, we first explored how CD3ζ ITAM mutations affect phosphorylation in the competitive inhibition model.

To test this effect, we used the median values of the best fit parameter sets from **Figure 4** and simulated LCK phosphorylation of CD3ζ with single or double ITAM mutations. The time course phosphorylation results for the competitive inhibition model are shown in **Supplemental Figure S5A**, while the results of the random order model are shown in **Supplemental Figure S5B**. In comparing these two figures, we see that there is a much smaller difference between the individual ITAMs in the competitive inhibition model compared to the random order model. This is particularly evident when comparing the results of ITAM C to ITAMs A and B in the single ITAM phosphorylation curves (xxC compared to Axx and xBx). These small differences in the half maximal time predicted by the competitive inhibition model are consistent with the data from (17).

Interestingly, while the competitive inhibition model does show a similar Hill coefficient between all of the CD3ζ ITAM mutant curves (**Supplemental Figure S5C**), there is a difference in the half maximal time, with fewer ITAMs resulting in a faster half maximal time (**Supplemental Figure S5D**). This is a similar effect shown by the experimental data from (17) in the presence of the phosphatase CD148. In this data, fewer ITAMs showed a higher EC50 for phosphatase inhibition. We, therefore, wanted to explore which mechanisms of phosphatase activity could allow the model to reproduce these results.

Keeping the same mechanism of LCK phosphorylation with competitive inhibition, we tried various mechanisms of dephosphorylation using parameters on the same order of magnitude as those for LCK phosphorylation. Specifically, we implemented a random order and phosphate dependent model for dephosphorylation. While a few parameter sets for the random order dephosphorylation mechanism or phosphate dependent dephosphorylation mechanism allowed for the Hill coefficients to be the same between all of the CD3ζ mutants, neither gave a clear increase in EC50 for increasing ITAM mutations.

On the other hand, with the competitive inhibition dephosphorylation mechanism, we were able to identify a defined parameter space that shows the same trends as seen in the experimental data for ITAM mutants in the presence of phosphatase inhibition (**Figure 5, Model BioNetGen Supplemental File S5, equations and parameters listed in Supplemental File S6**). This parameter space is characterized by phosphatase *K*_*M*_ values that are lower than those of LCK but follow the same trends in terms of the differences between tyrosine sites. A representative parameter set is shown in **Supplemental File S6**. This parameter space does not significantly depend on the catalytic rate of dephosphorylation (within an order of magnitude up or down from the baseline value) but does require that the competitive inhibition constant of the phosphatase (*X*_*l*_) be significantly less than one. Thus, we predict that CD3ζ is phosphorylated and dephosphorylated through a mechanism of competitive inhibition in which the phosphatase has a stronger binding preference for its substrates than LCK and more significant competitive inhibition.

**Figure 5:**
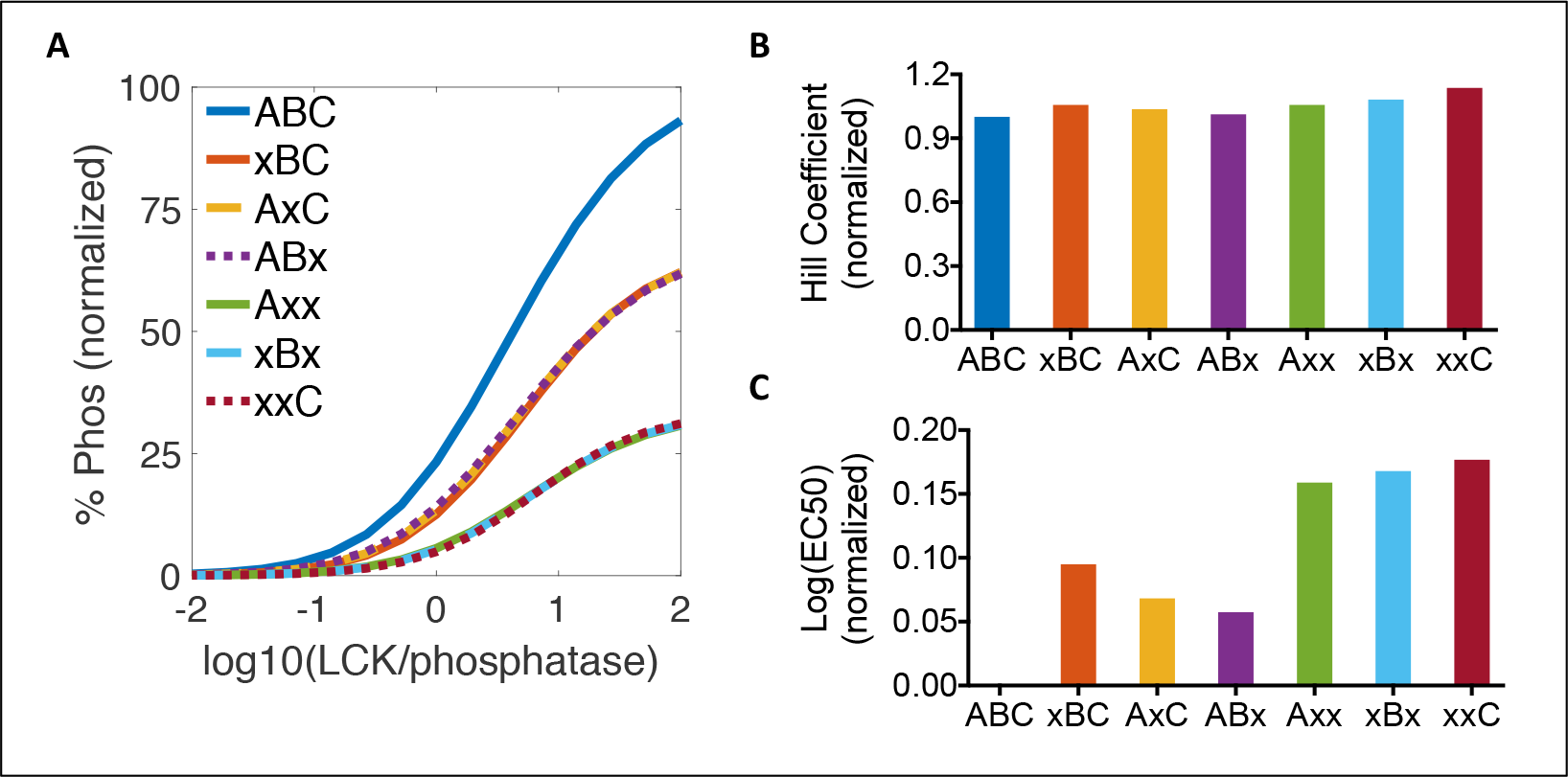
Model comparison to literature data of CD3ζ phosphorylation and dephosphorylation. (A) Predicted phosphorylation profiles for wild type, single, and double ITAM mutant CD3ζ. Mutated ITAMs are indicated by (x). The model was implemented using initial conditions described in the model from (17): 1 CD3z/μm^2^, 1000 LCK/μm^2^, and phosphatase concentrations between 10-100,000 molecules/μm^2^. (B) Hill coefficient of the predicted phosphorylation response for each CD3ζ mutant. (C) EC50 of the predicted phosphorylation response for each CD3ζ mutant.

### CD28 tyrosine sites are phosphorylated more slowly than CD3ζ tyrosine sites

Next, we investigated how the addition of a co-stimulatory domain, like CD28, could influence CAR phosphorylation. To test this, we inserted the intracellular domain of CD28 at the N-terminal of CD3ζ (28ζ), the same configuration typically used in the CAR constructs evaluated in pre-clinical studies and clinical trials (**Figure 6A**). CD28 has four tyrosine sites, each of which can be phosphorylated by LCK. We again used phospho-proteomic mass spectrometry to quantify the site-specific phosphorylation levels of 28ζ. **Supplemental Figure S6A** shows the sequence and trypsin cut sites of the CD28 intracellular domain, in which the second and third tyrosine sites in CD28 (Y206 and Y209) are both on the same peptide after trypsin digestion. Therefore, to individually measure the phosphorylation rates of these two sites, we made two more proteins with a tyrosine to phenylalanine mutation at each of these sites (28ζ-Y206F, and 28ζ-Y209F).

Interestingly, our measurements indicate that Y209 phosphorylation is required for the phosphorylation of Y206. **Supplemental Figure S6B** shows the individual CD28 tyrosine site phosphorylation time courses for each of the three CD28-CD3ζ recombinant proteins. From these graphs, we can see that there is no significant phosphorylation of the Y206 site without prior phosphorylation of Y209 (teal lines). In agreement with the literature (45), all tyrosine sites on CD28 are phosphorylated more slowly than the CD3ζ tyrosine sites. From the Y206F and Y209F mutants, we can see that mutating these sites reduces the overall phosphorylation rates of the CD28 protein, with almost no detectable CD28 phosphorylation in the Y209F mutant. This indicates that Y209, and to a lesser extent Y206, play a significant role in either the recruitment or phosphorylation activity of LCK toward CD28.

**Figure 6:**
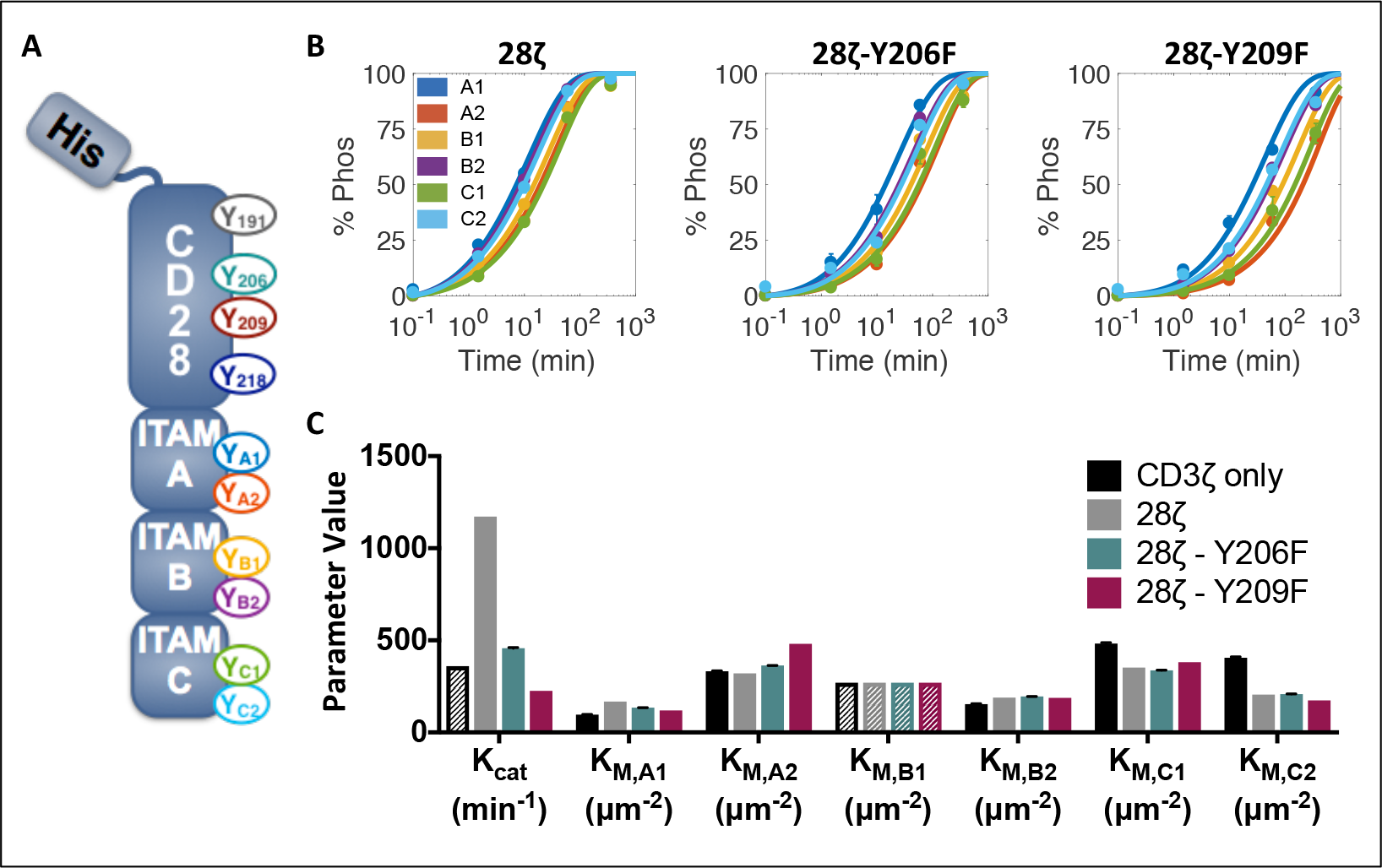
CD28 Influences CD3ζ phosphorylation kinetics. (A) Schematic of the His-tagged CD28-CD3ζ recombinant protein. (B) Experimental data (circles) and model fit (lines) for CD3z ITAM phosphorylation on wild type CD28-CD3ζ CD28-Y206F-CD3ζ, and CD28-Y209F-CD3ζ. Error bars represent the standard deviation of two technical replicates normalized by site-specific standard curves. (C) Estimated parameter sets. Solid bars show the mean and standard deviation of the 50 best fit parameter sets. Shaded bars show the value of parameters that were held constant during fitting.

### CD28 increases the phosphorylation rate of CD3ζ

CD28 influences the overall phosphorylation rate of CD3ζ, as well as the individual phosphorylation rate of site C2. **Figure 6B** (**dots**) shows the phosphorylation time courses of CD3ζ ITAM sites on the three CD28-CD3ζ recombinant proteins. Wild type CD28 increases the overall phosphorylation rate of all CD3ζ tyrosine sites, making it difficult to distinguish a specific order of phosphorylation in the 28ζ recombinant protein. To explore the mechanism of CD3ζ phosphorylation in the presence of CD28, we returned to our model of competitive inhibition. As each of the CD28 mutants show a clear change in the kinetics, we fit each of the data sets separately. In order to gain more mechanistic insight into the way that CD28 influences the phosphorylation, we attempted to independently fit the three types of parameters (i) catalytic rate, (ii) Michaelis-Menten constants, and (iii) inhibition constant scaling factor. None of the parameter types were able to provide an adequate fit on their own; however, fitting the catalytic rate and Michaelis-Menten constants together provided a good fit to the data (**Figure 6B, lines**). This fit was not improved by fitting all three parameters together or by fitting pair-wise combinations including the inhibition constant scaling factor.

The presence of CD28 results in a robust increase in the apparent catalytic rate of LCK phosphorylation (**Figure 6C**). This change in the catalytic rate is reduced for the CD28 mutants. Physiologically, this apparent catalytic rate likely represents a change in the local concentration of LCK due to recruitment by CD28. Comparing the changes in catalytic rate to those of the *K*_*M*_ values, we can see that the change in catalytic rate is much more significant, while there are only slight differences in the estimated *K*_*M*_ values. *K_M,C2_* shows the clearest difference, which accounts for its change in order of C2 now being phosphorylated before C1. This parameter fitting data confirms that the main mechanism of CD28 on CD3ζ is to increase the catalytic rate of LCK.

## Discussion

In this study, we used phospho-proteomic mass spectrometry and computational modeling to quantitatively assess the mechanism of CD3ζ and CD28 intracellular phosphorylation in a CAR construct. By measuring the phosphorylation of individual tyrosine sites on CD3ζ and CD28 over time, we showed that the six sites on CD3ζ and four sites on CD28 are phosphorylated at different rates. Individual CD3ζ point mutations showed that there is only a small amount of interaction between the CD3ζ ITAM sites, as removing one site does not greatly influence phosphorylation at any of the other sites. Additionally, attaching CD28 to the N-terminal of CD3ζ increased the overall phosphorylation rate of the protein, and particularly increased the relative rate of phosphorylation at the C2 ITAM site.

Interestingly, we did not see any effect of acidic lipid concentration on the phosphorylation rates of CD3ζ tyrosine sites. Previous studies have reported different effects of acidic lipids on CD3-family protein phosphorylation. Several studies have indicated that acidic lipids in the plasma membrane can control aberrant phosphorylation of CD3ε and CD28 in unstimulated cells through binding of basic residues in the protein to acidic lipids in the plasma membrane (39–41). In addition, Hui and Vale saw that ZAP-70 tandem SH2 domains were able to bind more quickly as CD3ζ became phosphorylated in a system containing 10% acidic POPS lipids compared to 0% POPS lipids (28). In our system, changing the concentration of acidic POPS lipids on the liposome surface (even up to 45% POPS) did not change the rate of phosphorylation of the protein as a whole or the relative phosphorylation rate order of the individual sites. Therefore, we believe that the acidic lipid concentration does not directly affect the phosphorylation of CD3ζ; but, as Hui and Vale showed, it may influence the binding kinetics of downstream proteins like ZAP-70 by more readily recruiting these proteins to membrane regions with acidic lipids.

We constructed a computational model to further investigate the mechanisms that lead to CD3ζ phosphorylation. With the model, we were able to identify a robust mechanism that accurately reflects the experimental data and calculate site-specific phosphorylation parameters, which are difficult to distinguish experimentally. Interestingly, previous work to attempt to determine an order for LCK phosphorylation of CD3ζ identified orders that differ from each other and from the order indicated by our measurements (16,46). These previous studies were performed in solution and with individual tyrosine peptides using techniques that limited their physiological relevance. However, in one study, the authors did use mass spectrometry to measure full length recombinant protein CD3ζ phosphorylation and found that, at intermediate time points, ITAM site A1 was significantly more phosphorylated than the other sites, which is consistent with our data (46).

To our knowledge, our work is the first study to specifically quantify the individual phosphorylation kinetics and phosphorylation mechanism of all six CD3ζ ITAM tyrosine sites on the same protein in a two-dimensional lipid-bound setting. We note that this recombinant protein system does make some modifications to the endogenous protein structure. In this study, we mutated out the CD3ζ tyrosine at site 64, which was shown to have no influence on the overall phosphorylation rate on the protein as a whole (**Supplemental Figure S2**). As such, it is unlikely that this mutation significantly changes the individual rates in such a way as to keep the total phosphorylation rate constant. One change to the protein that may play a more significant role is that the CAR proteins used in this liposomal system are not anchored by a transmembrane domain. Interactions within the extracellular or transmembrane domains of CAR proteins may play a role in the intracellular arrangement of CARs and their accessibility to LCK, thus influencing the phosphorylation rates. Despite these limitations, our work provides new mechanistic insights into the interplay among CD3ζ ITAM sites and between CD3ζ and CD28.

Our model provides novel insights into the effects of LCK-CD3ζ interactions. Our modeling results confirm a random order of LCK phosphorylation of CD3ζ, which validates previous studies in the literature based on average CD3ζ protein phosphorylation (17). The model also indicates that phosphorylated and unphosphorylated tyrosine sites on CD3ζ provide competitive feedback on one another and predicts that a similar mechanism is used by phosphatases in this system. This is significant as it provides an alternative mechanism to that of phosphate dependence, described by (17). Our competitive inhibition model is more robust and is able to reproduce a wider set of data. We believe that these two mechanisms are not entirely incompatible, as demonstrated by the modeling results including both of these effects together. Although, in this system, the effects of the phosphate dependent model were masked by the significantly stronger effects of competitive inhibition. Additionally, the insights from this modeling work could help inform other hypothesized models of LCK interaction with CD3ζ in the literature (14).

CD28 also plays an important role in the CAR structure, both by adding its co-stimulatory signaling and modulating the phosphorylation rates of CD3ζ. We showed that adding CD28 to the N-terminal of CD3ζ increases the overall phosphorylation of CD3ζ, and this is largely dependent on CD28-Y209. Our computational modeling work indicates that this is due to an increase in the effective catalytic rate of LCK. In the model, this parameter has the same effect as an increase in the local concentration of LCK. Importantly, our predictions agree with published experimental observations, where the Y206/Y209 region of CD28 has been implicated in the recruitment of LCK to the immunological synapse in endogenous T cells through binding of the SH2 domain on LCK (47). Here, we validate the importance of this site as a strong recruiter of LCK and its potential role in the strong activation of CD28-bearing CAR proteins.

Interestingly, CD28-Y209 is phosphorylated much more slowly than any of the sites on CD3ζ. In fact, when all of the CD3ζ sites are 100% phosphorylated, only about 25% of the CD28-Y209 sites are phosphorylated. This leads us to further hypothesize that unphosphorylated Y209 plays a role in recruiting LCK to the system. The CD28 Y206/Y209 sites are surrounded by multiple proline residues, and this proline-rich binding domain is likely responsible for LCK recruitment when CD28 Y206 and Y209 are unphosphorylated (36). A similar proline-rich region is also thought to help recruit LCK to CD3ε and CD2 (35,37). Perhaps this proline-rich Y206/Y209 site on CD28 is able to bind and recruit LCK more readily than other sites but, given its lower affinity for the catalytic pocket of LCK, it can be outcompeted by other sites on the same protein.

We also used our experimental system to explore the effects of the CD28-dependent reordering of CD3ζ ITAM phosphorylation. The data shows that CD28 increases the relative rate of site C2. This effect appears to be independent of phosphorylation at CD28 site Y209, indicating that another mechanism, such as the folding of the CD3ζ protein chain, must contribute to the increased phosphorylation at this particular site. More work needs to be done to decouple the binding preferences that lead to LCK recruitment from the catalytic activity of the protein-substrate pairs, to better understand how CD28 influences the relative order of CD3ζ phosphorylation.

Taken together, this work provides new insights into the activation of CAR-T cells through quantitative phospho-proteomic experiments and computational modeling. Our model predicts a single mechanism for LCK phosphorylation of CD3ζ ITAMs. In addition to producing novel measurements and a modeling framework that explains experimental observations, our work generates novel hypotheses regarding protein phosphorylation that can inform new experiments. In the future, this mechanistic insight about the CAR phosphorylation levels could be applied to better engineer CARs that are phosphorylated more quickly and to a greater extent, and hence, more optimally activate T cells for therapeutic purposes.

## Author Contributions

J.A.R., P.W. and S.D.F. conceived of and designed the project. J.A.R. performed the experiments, advised by P.W. D.Z. acquired the phospho-proteomic data, advised by N.A.G. J.A.R. analyzed the proteomic data, advised by N.A.G. J.A.R performed the computational modeling, advised by S.D.F. J.A.R. wrote the original draft. All co-authors contributed to reviewing and editing the manuscript. J.A.R, N.A.G, P.W. and S.D.F. provided funding and resources to support the project.

## Acknowledgements

The authors would like to thank the lab of Ronald Vale at UCSF for providing the recombinant protein plasmids. This work was supported by the National Cancer Institute of the National Institutes of Health under Award Numbers F31CA200242 (to J.A.R.), R01EB017206, R01CA170820, and P01CA132681 (to P.W.)

## Conflicts of Interest

The authors have no conflicts of interest to disclose.

## Supplemental Information

**Figure S1**: **Standard curves for phosphorylated:unphosphorylated peptide intensity normalization**.

Data was collected on the mass spectrometer in three sets, and each set was normalized by a standard curve collected at the same time. All of the standard curves used are shown here, with each row being used to analyze a separate set of data. Row 1 was used to analyze the first biological replicate of the wild type ITAM phosphorylation on 10% POPS liposomes and the wild type ITAM phosphorylation on 0% and 45% POPS liposomes. Row 2 was used to analyze the second biological replicate of the wild type ITAM phosphorylation on 10% POPS liposomes as well as the individual tyrosine to phenylalanine CD3ζ ITAM point mutants. The third and fourth rows were used to analyze all 28ζ proteins, including the Y206F and Y209F mutants.

**Figure S2**: **Comparison of wild type and Y64F CD3ζ phosphorylation rates**.

(A, B) Western blot of CD3ζ phosphorylation time courses. Liposomes bearing ~10,000 molecules/μm^2^ of wild type CD3ζ (A) or CD3ζ-Y64F (B) and ~155 molecules/μm^2^ of LCK-Y505F were allowed to react in the presence of ATP, as described in the Methods. At various times, the reaction was stopped by adding one volume of 10x SDS PAGE running buffer (Millipore Sigma) and boiling for 5 minutes. Samples were then analyzed by western blotting with anti-phospho-tyrosine antibodies. Blots were analyzed using the C-DiGit Western Blot Scanner (Li-Cor Biosciences - U.S.). (C) Quantification of CD3ζ phospho-tyrosine western blots. The intensity of the CD3ζ monomer bands (~23 kDa) was analyzed using the Image Studio Digits Software (Li-Cor Biosciences - U.S., version 3.1). Signal intensities for each protein (wild type and Y64F mutant) were normalized by the signal at 60 minutes and a standard sigmoidal curve was fit to the data.

**Figure S3**: **Statistical comparison of CD3ζ ITAM site phosphorylation levels**.

(Blue) 10 minute comparison, (Orange) 60 minute comparison. Measured by pairwise Tukey t-tests (**** p<0.0001, *** p<0.001, ** p<0.01, * p<0.05, n.s. not significant).

**Figure S4: CD3ζ ITAM site phosphorylation model fits**.

(A) The second biological replicate of wild type CD3ζ ITAM protein stimulated on liposomes containing 10% POPS lipids, (B) wild type CD3ζ ITAM protein stimulated on liposomes containing 0% POPS lipids, (C) wild type CD3ζ ITAM protein stimulated on liposomes containing 45% POPS lipids. (D-J) CD3ζ proteins bearing individual tyrosine to phenylalanine ITAM point mutations stimulated on liposomes containing 10% POPS lipids.

**Figure S5**: **Model predictions for ITAM mutant phosphorylation profiles**.

(A) Predicted phosphorylation profiles for wild type, single, and double ITAM mutant CD3ζ with the competitive inhibition model. Mutated ITAMs are indicated by (x). Model was implemented using the optimal parameters from the model fitting shown in **Figure 3D** with initial conditions of 20 LCK/μm^2^ and 2000 CD3z/μm^2^.

(B) Predicted phosphorylation profiles for wild type, single, and double ITAM mutant CD3ζ with the random order model. Mutated ITAMs are indicated by (x). Model was implemented using the optimal parameters from the model fitting shown in **Figure 3B** with initial conditions of 20 LCK/μm^2^ and 2000 CD3ζ/μm^2^.

(C) Hill coefficient of the phosphorylation response predicted by the model for each CD3ζ mutant in the competitive inhibition model.

(D) Half maximal time of the predicted phosphorylation response for each CD3ζ mutant in the competitive inhibition model.

**Figure S6**: **CD28 phosphorylation profiles**.

(A) Sequence of CD28 intracellular domain with trypsin cut sites denoted. Individual tyrosine sites are labeled in different colors.

(B) Experimental data for CD28 tyrosine site phosphorylation on wild type CD28-CD3ζ, CD28-Y206F-CD3ζ, and CD28-Y209F-CD3ζ. Error bars represent the standard deviation of two technical replicates normalized by site-specific standard curves.

**File S1**: RandomOrder.bngl BioNetGen file of CD3ζ LCK sequential order phosphorylation model. Mean optimal parameter sets are listed in this file.

**File S2**: SequentialOrder.bngl BioNetGen file of CD3ζ LCK random order phosphorylation model. Mean optimal parameter sets are listed in this file.

**File S3**: PhosphateDependent.bngl BioNetGen file of CD3ζ LCK phosphate dependent phosphorylation model. Mean optimal parameter sets are listed in this file.

**File S4**: CompetitiveInhibition.bngl BioNetGen file of CD3ζ LCK competitive inhibition phosphorylation model. Mean optimal parameter sets are listed in this file.

**File S5**: CompetitiveInhibition_phosphatase.bngl BioNetGen file of CD3ζ LCK phosphatase competitive inhibition model. Mean optimal parameter sets are listed in this file.

**File S6**: LCK-phosphatase competitive inhibition model. List of the model parameters and equations used to simulate the competitive inhibition mechanism of LCK and phosphatase interaction with CD3ζ.

## References

1. Fesnak AD, June CH, Levine BL. Engineered T cells: The promise and challenges of cancer immunotherapy. Nat Rev Cancer [Internet]. 2016;16(9):566–81. Available from: http://dx.doi.org/10.1038/nrc.2016.97

2. Firor AE, Jares A, Ma Y. From humble beginnings to success in the clinic: Chimeric antigen receptor-modified T-cells and implications for immunotherapy. Exp Biol Med. 2015;240(8):1087–98.

3. Mullard A. Second anticancer CAR T therapy receives FDA approval. Nat Rev Drug Discov [Internet]. 2017;16(12):818–818. Available from: http://www.nature.com/doifinder/10.1038/nrd.2017.249

4. Kingwell K. CAR T therapies drive into new terrain. Nat Rev Drug Discov [Internet]. 2017;16(5):301–4. Available from: http://dx.doi.org/10.1038/nrd.2017.84

5. Morgan RA, Yang JC, Kitano M, Dudley ME, Laurencot CM, Rosenberg SA. Case report of a serious adverse event following the administration of t cells transduced with a chimeric antigen receptor recognizing ERBB2. Mol Ther [Internet]. 2010;18(4):843–51. Available from: http://dx.doi.org/10.1038/mt.2010.24

6. Lee DW, Kochenderfer JN, Stetler-Stevenson M, Cui YK, Delbrook C, Feldman SA, et al. T cells expressing CD19 chimeric antigen receptors for acute lymphoblastic leukaemia in children and young adults: A phase 1 dose-escalation trial. Lancet. 2015;385(9967):517–28.

7. Moon EK, Wang LC, Dolfi D V., Wilson CB, Ranganathan R, Sun J, et al. Multifactorial T-cell hypofunction that is reversible can limit the efficacy of chimeric antigen receptor-transduced human T cells in solid tumors. Clin Cancer Res. 2014;20(16):4262–73.

8. Harris DT, Hager M V, Smith SN, Stone JD, Kruger P, Lever M, et al. Comparison of T Cell Activities Mediated by Human TCRs and CARs That Use the Same Recognition Domains. J Immunol. 2018;

9. Palacios EH, Weiss A. Function of the Src-family kinases, Lck and Fyn, in T-cell development and activation. Oncogene. 2004;23(48 REV. ISS. 7):7990–8000.

10. Lovatt M, Filby A, Parravicini V, Werlen G, Palmer E, Zamoyska R. Lck Regulates the Threshold of Activation in Primary T Cells, While both Lck and Fyn Contribute to the Magnitude of the Extracellular Signal-Related Kinase Response. Mol Cell Biol [Internet]. 2006;26(22):8655–65. Available from: http://mcb.asm.org/cgi/doi/10.1128/MCB.00168-06

11. Nika K, Soldani C, Salek M, Paster W, Gray A, Etzensperger R, et al. Constitutively active Lck kinase in T cells drives antigen receptor signal transduction. Immunity [Internet]. 2010 Jun 25 [cited 2014 Mar 17];32(6):766–77. Available from: http://www.pubmedcentral.nih.gov/articlerender.fcgi?artid=2996607&tool=pmcentrez&rendertype=abstract

12. Chakraborty AK, Weiss A. Insights into the initiation of TCR signaling. Nat Immunol [Internet. 2014 Sep 19 [cited 2015 Jul 29];15Chakrabo(9):798–807. Available from: http://www.nature.com/ni/journal/v15/n9/abs/ni.2940.html#affil-auth

13. Au-Yeung BB, Deindl S, Hsu L-Y, Palacios EH, Levin SE, Kuriyan J, et al. The structure, regulation, and function of ZAP-70. Immunol Rev [Internet]. 2009 Mar;228(1):41–57. Available from: http://www.ncbi.nlm.nih.gov/pubmed/19290920

14. Sjolin-goodfellow H, Frushicheva MP, Ji Q, Cheng DA, Kadlecek TA, Cantor AJ, et al. The catalytic activity of the kinase ZAP-70 mediates basal signaling and negative feedback of the T cell receptor pathway. Sci Signal. 2015;8(377):1–14.

15. Osman N, Turner H, Lucas S, Reif K, Cantrell DA. The protein interactions of the immunoglobulin receptor family tyrosine-based activation motifs present in the T cell receptor zeta subunits and the CD3 gamma, delta and epsilon chains. Eur J Immunol [Internet]. 1996;26(5):1063–8. Available from: http://www.ncbi.nlm.nih.gov/entrez/query.fcgi?cmd=Retrieve…db=PubMed…dopt=Citation…list_uids=8647168%5Cnhttp://onlinelibrary.wiley.com/store/10.1002/eji.1830260516/asset/1830260516_ftp.pdf?v=1…t=hu5vxjw1…s=0206b992dacd709a851e9c846c90969abec002ef

16. Kersh EN. Fidelity of T Cell Activation Through Multistep T Cell Receptor Phosphorylation. Science (80-) [Internet]. 1998 Jul 24 [cited 2014 Mar 8];281(5376):572–5. Available from: http://www.sciencemag.org/cgi/doi/10.1126/science.281.5376.572

17. Mukhopadhyay H, Wet B De, Clemens L, Maini PK, Allard J, van der Merwe PA, et al. Multisite phosphorylation of the T cell receptor Z-chain modulates potency but not the switch-like response. Biophys J. 2016;110(8):1896–906.

18. Mukhopadhyay H, Cordoba S-P, Maini PK, van der Merwe PA, Dushek O. Systems model of T cell receptor proximal signaling reveals emergent ultrasensitivity. PLoS Comput Biol [Internet]. 2013 Jan [cited 2014 Jan 22];9(3):e1003004. Available from: http://www.pubmedcentral.nih.gov/articlerender.fcgi?artid=3610635&tool=pmcentrez&rendertype=abstract

19. Boomer JS, Green JM. An enigmatic tail of CD28 signaling. Cold Spring Harb Perspect Biol [Internet]. 2010 Aug [cited 2014 Apr 11];2(8):a002436. Available from: http://www.pubmedcentral.nih.gov/articlerender.fcgi?artid=2908766&tool=pmcentrez&rendertype=abstract

20. Holdorf AD, Lee K-H, Burack WR, Allen PM, Shaw AS. Regulation of Lck activity by CD4 and CD28 in the immunological synapse. Nat Immunol [Internet]. 2002 Mar [cited 2014 Jan 27];3(3):259–64. Available from: http://www.ncbi.nlm.nih.gov/pubmed/11828322

21. Carey KD, Dillon TJ, Schmitt JM, Baird a M, Holdorf a D, Straus DB, et al. CD28 and the tyrosine kinase lck stimulate mitogen-activated protein kinase activity in T cells via inhibition of the small G protein Rap1. Mol Cell Biol. 2000;20(22):8409–19.

22. Tian R, Wang H, Gish GD, Petsalaki E, Pasculescu A, Shi Y, et al. Combinatorial proteomic analysis of intercellular signaling applied to the CD28 T-cell costimulatory receptor. Proc Natl Acad Sci U S A [Internet]. 2015;112(13):E1594–603. Available from: http://www.ncbi.nlm.nih.gov/pubmed/25829543

23. Teng JM, King PD, Sadra A, Liu X, Han A, Selvakumar A, et al. Phosphorylation of each of the distal three tyrosines of the CD28 cytoplasmic tail is required for CD28-induced T cell IL-2 secretion. Tissue Antigens [Internet]. 1996;48(4 Pt 1):255–64. Available from: http://www.ncbi.nlm.nih.gov/entrez/query.fcgi?cmd=Retrieve&db=PubMed&dopt=Citation&list_uids=8946678

24. Lu Y, Phillips C a, Bjorndahl JM, Trevillyan JM. CD28 signal transduction: tyrosine phosphorylation and receptor association of phosphoinositide-3 kinase correlate with Ca(2+)-independent costimulatory activity. Eur J Immunol [Internet]. 1994;24(11):2732–9. Available from: http://www.ncbi.nlm.nih.gov/pubmed/7957566

25. Michel F, Attal-Bonnefoy G, Mangino G, Mise-Omata S, Acuto O. CD28 as a molecular amplifier extending TCR ligation and signaling capabilities. Immunity [Internet]. 2001 Dec;15(6):935–45. Available from: http://www.ncbi.nlm.nih.gov/pubmed/11754815

26. Chae W-J, Lee H-K, Han J-H, Kim S-WV, Bothwell ALM, Morio T, et al. Qualitatively differential regulation of T cell activation and apoptosis by T cell receptor zeta chain ITAMs and their tyrosine residues. Int Immunol. 2004; 16(9):1225–36.

27. Kersh BEN, Kersh GJ, Allen PM. Partially Phosphorylated T Cell Receptor Zeta Molecules Can Inhibit Inhibit T Cell Activation. J Exp Med. 1999;190(11).

28. Hui E, Vale RD. In vitro membrane reconstitution of the T-cell receptor proximal signaling network. Nat Struct Mol Biol [Internet]. 2014 Feb [cited 2014 Mar 21];21(2):133–42. Available from: http://www.ncbi.nlm.nih.gov/pubmed/24463463

29. Rohrs JA, Wang P, Finley SD. Predictive Model of Lymphocyte-Specific Protein Tyrosine Kinase (LCK) Autoregulation. Cell Mol Bioeng. 2016;9(3):351–67.

30. Li H, Korennykh a V, Behrman SL, Walter P. Mammalian endoplasmic reticulum stress sensor IRE1 signals by dynamic clustering. Proc Natl Acad Sci U S A [Internet]. 2010;107(37):16113–8. Available from: http://www.ncbi.nlm.nih.gov/pubmed/20798350

31. Pear WS, Nolan GP, Scott ML, Baltimore D. Production of high-titer helper-free retroviruses by transient transfection (retroviral packaing cells/gene therapy). Cell Biol. 1993;90(September):8392–6.

32. Faeder JR, Blinov ML, Hlavacek WS. Rule-based modeling of biochemical systems with BioNetGen. Methods Mol Biol. 2009;500(1):113–67.

33. Iadevaia S, Lu Y, Morales FC, Mills GB, Ram PT. Identification of optimal drug combinations targeting cellular networks: integrating phospho-proteomics and computational network analysis. Cancer Res [Internet]. 2010 Sep 1 [cited 2014 Nov 12];70(17):6704–14. Available from: http://www.pubmedcentral.nih.gov/articlerender.fcgi?artid=2932856&tool=pmcentrez&rendertype=abstract

34. Lee KA, Craven KB, Niemi GA, Hurley JB. Mass spectrometric analysis of the kinetics of in vivo rhodopsin phosphorylation. Protein Sci [Internet]. 2002;11(4):862–74. Available from: http://www.ncbi.nlm.nih.gov/pubmed/11910029%5Cnhttp://onlinelibrary.wiley.com/store/10.1110/ps.3870102/asset/110862_ftp.pdf?v=1&t=i7vm0sqx&s=342f456a0d9137f5b52b88a9dd0041bb3b7b8668

35. Bell GM, Fargnoli J, Bolen JB, Kish L, Imboden JB. The SH3 domain of p56lck binds to proline-rich sequences in the cytoplasmic domain of CD2. J Exp Med [Internet]. 1996;183(1):169–78. Available from: http://www.pubmedcentral.nih.gov/articlerender.fcgi?artid=2192399&tool=pmcentrez&rendertype=abstract

36. Holdorf AD, Green JM, Levin SD, Denny MF, Straus DB, Link V, et al. Proline residues in CD28 and the Src homology (SH)3 domain of Lck are required for T cell costimulation. J Exp Med [Internet]. 1999;190(3):375–84. Available from: http://www.ncbi.nlm.nih.gov/pubmed/10430626%5Cnhttp://www.pubmedcentral.nih.gov/articlerender.fcgi?artid=PMC2195584

37. Li L, Guo X, Shi X, Li C, Wu W, Yan C, et al. Ionic CD3-Lck interaction regulates the initiation of T-cell receptor signaling. Proc Natl Acad Sci [Internet]. 2017;114(29):E5891–9. Available from: http://www.pnas.org/lookup/doi/10.1073/pnas.1701990114

38. Lee TY, Hsu JBK, Chang WC, Huang H Da. RegPhos: A system to explore the protein kinase-substrate phosphorylation network in humans. Nucleic Acids Res. 2011;39(SUPPL. 1):777–87.

39. Aivazian D, Stern LJ. Phosphorylation of T cell receptor zeta is regulated by a lipid dependent folding transition. Nat Struct Biol [Internet]. 2000 Nov;7(11):1023–6. Available from: http://www.ncbi.nlm.nih.gov/pubmed/11062556

40. Dobbins J, Gagnon E, Godec J, Pyrdol J, Vignali DAA, Sharpe AH, et al. Binding of the cytoplasmic domain of CD28 to the plasma membrane inhibits Lck recruitment and signaling. Sci Signal. 2016;9(438):1–13.

41. Gagnon E, Schubert D a, Gordo S, Chu HH, Wucherpfennig KW. Local changes in lipid environment of TCR microclusters regulate membrane binding by the CD3ε cytoplasmic domain. J Exp Med [Internet]. 2012;209(13):2423–39. Available from: http://www.pubmedcentral.nih.gov/articlerender.fcgi?artid=3526357&tool=pmcentrez&rendertype=abstract

42. Salazar C, Höfer T. Versatile regulation of multisite protein phosphorylation by the order of phosphate processing and protein-protein interactions. FEBS J. 2007;274(4):1046–61.

43. Ferrell JE. Tripping the switch fantastic: How a protein kinase cascade can convert graded inputs into switch-like outputs. Trends Biochem Sci. 1996;21(12):460–6.

44. Ferrell JE, Ha SH. Ultrasensitivity part II□: multisite phosphorylation, stoichiometric inhibitors, and positive feedback. 2014;39(11).

45. Hui E, Cheung J, Zhu J, Su X, Taylor MJ, Wallweber HA, et al. T cell costimulatory receptor CD28 is a primary target for PD-1-mediated inhibition. Science (80-) [Internet]. 2017;4(March):eaaf1292. Available from: http://www.sciencemag.org/lookup/doi/10.1126/science.aaf1292

46. Housden HR, Skipp PJS, Crump MP, Broadbridge RJ, Crabbe T, Perry MJ, et al. Investigation of the kinetics and order of tyrosine phosphorylation in the T-cell receptor zeta chain by the protein tyrosine kinase Lck. Eur J Biochem. 2003;270:2369–76.

47. Hofinger E, Sticht H. Multiple modes of interaction between Lck and CD28. J Immunol [Internet]. 2005;174(7):3839–3840; author reply 3840. Available from: http://www.jimmunol.org/content/174/7/3839.2.full

